# Position information encoded by population activity in hierarchical visual areas

**DOI:** 10.1101/073940

**Authors:** Kei Majima, Paul Sukhanov, Tomoyasu Horikawa, Yukiyasu Kamitani

## Abstract

Neurons in high-level visual areas respond to more complex visual features with broader receptive fields (RFs) compared to those in low-level visual areas. Thus, high-level visual areas are generally considered to carry less information regarding the position of seen objects in the visual field. However, larger RFs may not imply loss of position information at the population level. Here, we evaluated how accurately the position of a seen object could be predicted (decoded) from activity patterns in each of six representative visual areas with different RF sizes (V1–V4, LOC, and FFA). We collected fMRI responses while human subjects viewed a ball randomly moving in a two-dimensional field. To estimate population RF sizes of individual fMRI voxels, RF models were fitted for individual voxels in each brain area. The voxels in higher visual areas showed larger estimated RFs than those in lower visual areas. Then, the ball’s position in a separate session was predicted by maximum likelihood regression (support vector regression, SVR) to predict the position. We found that regardless of the difference in RF size, all visual areas showed similar prediction accuracies, especially on the horizontal dimension. Higher areas showed slightly lower accuracies on the vertical dimension, which appears to be attributed to the narrower spatial distributions of the RFs centers. The results suggest that much of position information is preserved in population activity through the hierarchical visual pathway regardless of RF sizes, and is potentially available in later processing for recognition and behavior.

**Significance statement:** High-level ventral visual areas are thought to achieve position invariance with larger receptive fields at the cost of the loss of precise position information. However, larger receptive fields may not imply loss of position information at the population level. Here, multivoxel fMRI decoding reveals that high-level visual areas are predictive of an object’s position with similar accuracies to low-level visual areas, especially on the horizontal dimension, preserving the information potentially available for later processing.

## Introduction

Along the ventral visual cortical pathway, neurons in higher-level areas respond to more complex visual features with broader receptive fields (RFs). This is thought to serve to represent objects regardless of the position in the visual field. Because of this receptive field property, position information is often assumed to be lost in these areas (Ito et al., 1995; Logothetis and Sheinberg, 1996; Tanaka, 1996). However, the loss of position information in single neurons does not necessarily imply the loss of position information at the population level. Theoretical studies have suggested that if the RFs of model neurons are uniformly distributed in the 2D visual field, the Fisher information about the position of a stimulus is not degraded by an increase in RF size (Zhang and Sejnowski, 1999; Eurich and Wilke, 2000). As the Fisher information provides the theoretical lower bound of the estimation/decoding error, position information may not be lost even in visual areas with large RFs, such as the lateral occipital complex (LOC) and fusiform face area (FFA). While several recent fMRI studies demonstrated successful classification of the position (e.g. left vs. right, upper vs. lower) of a presented object from ventral visual areas (Schwarzlose et al., 2008; Carlson et al., 2011; Golomb and Kanwisher, 2011; Kay et al., 2015), the relationship between RF size and decoded position information across visual areas has not been quantitatively examined.

Here, we estimated RF sizes for fMRI voxels and evaluated how accurately the position of a seen object was predicted (decoded) from activity patterns in each of six representative visual areas (V1–V4, LOC, and FFA). In our experiments, we collected fMRI responses while subjects viewed a ball randomly moving in a two-dimensional field (Figure 1; a ball with a diameter of 1.6° presented within a 7.6° × 7.6° square field). The subjects were instructed to fix their eyes to the fixation point and keep track of the ball in their mind. fMRI activity was collected at a 3 × 3 × 3 mm resolution, and the signals from voxels in areas V1–V4, LOC and FFA were analyzed (see Materials and methods). To estimate RF sizes, RF models were fitted for individual voxels in each brain area (Dumoulin and Wandell, 2008). In the decoding analysis, the ball position was predicted either by maximum likelihood estimation using the RF models of individual voxels or by support vector regression (SVR; Drucker et al., 1997; Chang and Lin, 2011) with multivoxel patterns as inputs (Figure 1; see Materials and methods). While the maximum likelihood method provides straightforward interpretation given accurate RF models, SVR is expected to perform model-free information retrieval from fMRI data. Our results show that with both methods, position decoding accuracies were similar across the low- and high-level visual areas, especially along the horizontal axis, despite the differences in RF size and spatial coding in individual voxels.

**Figure 1.**
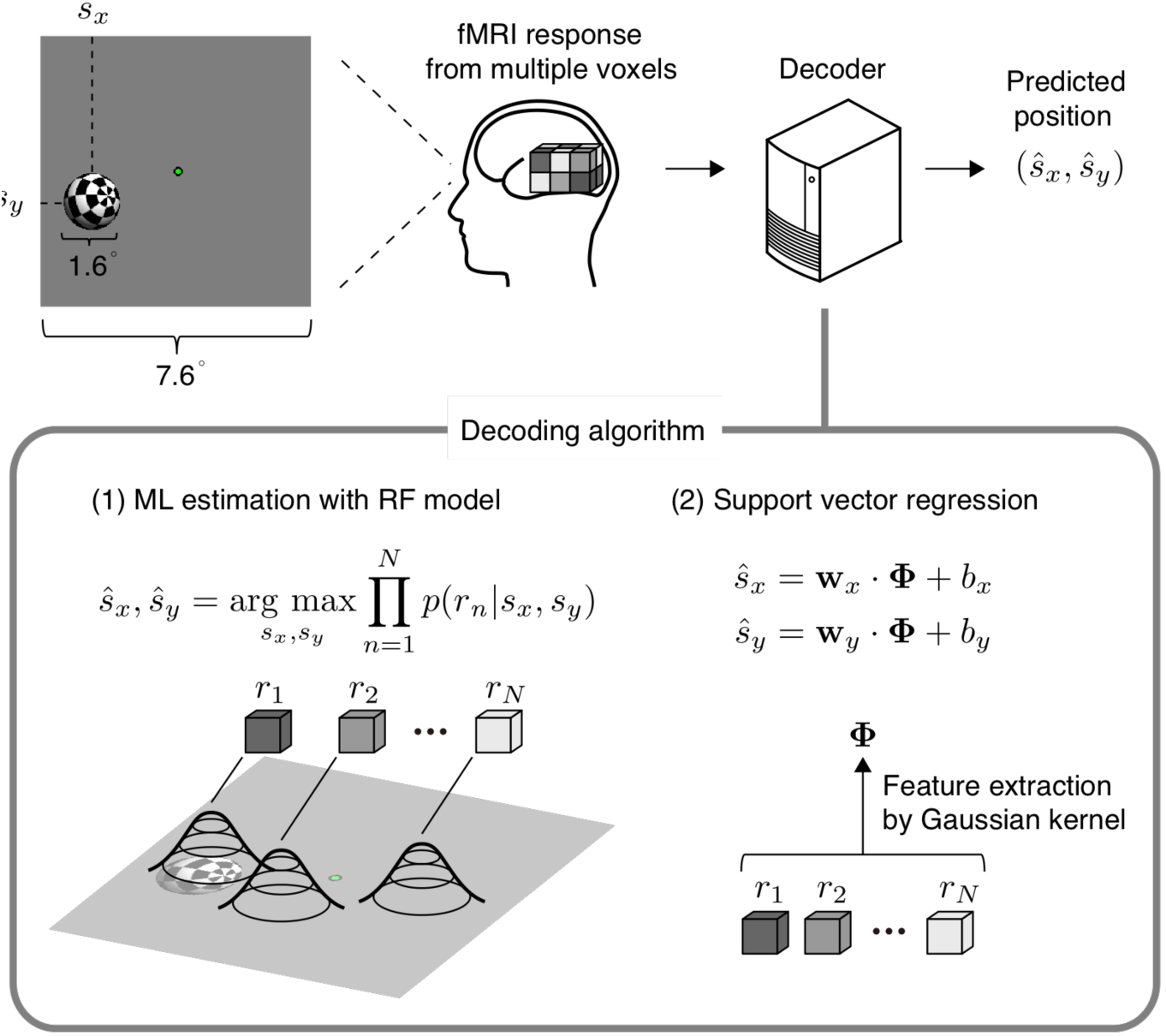
Visual stimulus and analysis overview. A white-and-black checkered sphere was displayed on a screen with a flickering rate of 6 Hz. The notations show the size of the sphere and the size of the field where the sphere could move. The position of the center of the sphere was predicted from measured brain activity. Prediction was performed based on maximum likelihood estimation using estimated receptive field models or the support vector regression algorithm.

## Materials and methods

### Subjects

Five healthy subjects (one female and four males, aged between 23 and 31) with normal or corrected-to-normal vision participated in our experiments. This sample size was chosen based on previous fMRI studies with similar experimental designs (Dumoulin and Wandell, 2008; Amano et al., 2009). We obtained written informed consent from all subjects prior to their participation in the experiments, and the Ethics Committee of ATR approved the study protocol.

### Position tracking experiment

The stimulus was created with Psychtoolbox-3 (http://psychtoolbox.org/)(RRID: SCR_002881) and the associated openGL for Psychtoolbox extension. The stimulus was projected onto a display in the fMRI scanner and viewed through a mirror attached to the headcoil. We conducted three scanning sessions (runs) for each subject. In each run, an initial rest period of 32 s was followed by four blocks of stimulus presentation, which each lasted for 240 s. The stimulus presentation blocks were separated by 12-s rest periods. An extra 12-s rest period was added to the end of each run (1,040 s total for each run). During each of the rest periods, a circular fixation point (0.25° diameter) was displayed on the center of the display and subjects were instructed to attend to this point. During stimulus presentation, in addition to the fixation point, a white-and-black checkered sphere with a diameter of 1.6° was displayed with a flickering rate of 6 Hz (Figure 1).

The sphere was programmed to move in a random orbit produced by the following process. For each frame, the position of the center of the sphere was updated by

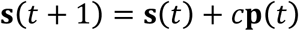

where ***s***(*t*) is the position at frame *t* (i.e. ***s***(*t*) *=* (*s_x_*(*t*), *s_y_*(*t*))) and **p**(*t*) is the vector indicating the direction of the movement from frame *t* to (*t +* 1), which imitates momentum. The constant *c,* which is a parameter that controls the speed, was set to 0.008 in this study. The vector **p**(*t*) was updated by

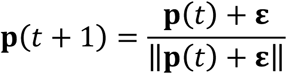

where **ε** is a random vector sampled from a two dimensional Gaussian distribution *𝒩*(0*,σ^2^I*) for every frame. *σ* was set to 0.1 in this study. The movement of the sphere center was limited within a 6.0° × 6.0° square field (the stimulus spanned a 7.6° × 7.6° square field. We restricted the stimulus position within this range so that subjects could easily keep track of the target sphere with attention. If ***s***(*t +* 1) was not in the allowed region in terms of horizontal or vertical position, the first or second element of **p**(*t*) was multiplied by −1 before the position was updated. This procedure ensures that the sphere is bound to the edge of the allowed region. The frame rate of stimulus presentation was 60 Hz.

### Retinotopy experiment

The retinotopy experiments were conducted according to the conventional protocol (Engel et al., 1994; Sereno et al., 1995). We used a rotating wedge and an expanding ring covered in a flickering checkerboard. The data were used to delineate the borders between visual cortical areas, and to identify the retinotopic map (V1–V4) on the flattened cortical surfaces of individual subjects.

### Localizer experiment

The functional localizer experiments were conducted to identify the lateral occipital complex (LOC)(Kourtzi and Kanwisher, 2000) and fusiform face area (FFA)(Kanwisher et al., 1997) for each individual subject. The localizer experiments comprised four to eight runs, and each run contained 16 stimulus blocks. In the experiments, intact or scrambled images (12° × 12°) belonging to face, object, house, and scene categories were presented around the center of the screen. Stimuli from each of the eight stimulus types (four categories × two conditions) were presented twice per run. Each stimulus block consisted of a 15-s intact or scrambled stimulus presentation. The intact and scrambled stimulus blocks were presented successively (the order of the intact and scrambled stimulus blocks was random), followed by a 15-s rest period where a uniform gray background was displayed. Extra 33-s and 6-s rest periods were presented before and after each run, respectively. In each stimulus block, 20 different images of the same stimulus type were presented for 0.3 s, separated by 0.4-second-long blank intervals.

### MRI acquisition

We collected fMRI data using a 3.0-Tesla Siemens MAGNETOM Trio a Tim scanner located at the ATR Brain Activity Imaging Center. An interleaved T2*-weighted gradient-EPI scan was performed to acquire functional images of the entire occipital lobe (position tracking experiment and retinotopy experiment: TR, 2,000 ms; TE, 30 ms; flip angle, 80 deg; FOV, 192 × 192 mm; voxel size, 3 × 3 × 3 mm; slice gap, 0 mm; number of slices, 30; localizer experiment: TR, 3,000 ms; TE, 30 ms; flip angle, 80 deg; FOV, 192 × 192 mm; voxel size, 3 × 3 × 3 mm; slice gap, 0 mm; number of slices, 50). T2-weighted turbo spin echo images were scanned to acquire high-resolution anatomical images of the same slices used for the EPI (position tracking experiment and retinotopy experiment: TR, 6,000 ms; TE, 57 ms; flip angle, 160 deg; FOV, 192 × 192 mm; voxel size, 0.75 × 0.75 × 3.0 mm; localizer experiment: TR, 7,020 ms; TE, 69 ms; flip angle, 160 deg; FOV, 192 × 192 mm; voxel size, 0.75 × 0.75 × 3.0 mm). T1-weighted magnetization-prepared rapid acquisition gradient-echo (MP-RAGE) fine-structural images of the entire head were also acquired (TR, 2,250 ms; TE, 3.06 ms; TI, 900 ms; flip angle, 9 deg, FOV, 256 × 256 mm; voxel size, 1.0 × 1.0 × 1.0 mm).

### MRI data preprocessing

The first 8-s scans (position tracking experiment and retinotopy experiment) or 9-s scans (localizer experiment) of each run were discarded to avoid MRI scanner instability. We then subjected the acquired fMRI data to three-dimensional motion correction with SPM5 (http://www.fil.ion.ucl.ac.uk/spm). Those data were then coregistered to the within-session high-resolution anatomical images of the same slices used for EPI and subsequently to the whole-head high-resolution anatomical images. The coregistered data were then re-interpolated as 3 × 3 × 3 mm voxels. For the data from the position tracking experiment, the signal amplitudes from individual voxels were linearly detrended in each run and shifted by 4 s (two fMRI volumes) to compensate for hemodynamic delay. We did not perform the convolution with the stimulus with a hemodynamic response function, as it makes the estimation of receptive field models difficult. Instead, we moved the target very slowly so that each fMRI volume can be associated with a single position (with a 4-s delay).

### Region of interest (ROI) selection

V1, V2, V3, and V4 were identified using the data from the retinotopy experiments (Engel et al., 1994; Sereno et al., 1995). The lateral occipital complex (LOC) and fusiform face area (FFA) were identified using the data from the functional localizer experiments (Kanwisher et al., 1997; Kourtzi and Kanwisher, 2000). The data from the retinotopy experiment were transformed into Talairach space and the visual cortical borders were delineated on the flattened cortical surfaces using BrainVoyager QX (http://www.brainvoyager.com)(RRID: SCR_013057). The coordinates of voxels around the gray-white matter boundary in V1–V4 were identified and transformed back into the original coordinates of the EPI images. The localizer experiment data were analyzed using SPM5. The voxels showing significantly higher activation in response to intact object or face images compared with that for scrambled images (*t*-test, uncorrected *p* < 0.05 or 0.01) were identified, and defined as LOC and FFA, respectively.

### Population receptive field model fitting

To estimate the receptive field, we fitted a population receptive field model to voxel amplitudes from each voxel in the visual cortex. We used fMRI data from the position tracking experiment in the analysis. Our model was based on a two-dimensional Gaussian receptive field and the noise on voxel amplitudes was assumed to be Gaussian (Dumoulin and Wandell, 2008). Mathematically, this assumption was expressed by

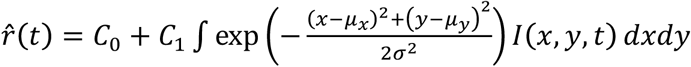

and

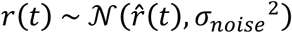

*r*̂ (*t*) and *r*(*t*) are the fitted and observed voxel amplitudes for the *t*-th fMRI volume. *C*_0_, *C*_1_, *μ_x_*, *μ_y_*, *σ*, and *σ_noise_* are constants to be estimated. *I*(*x*,*y*, *t*) is the binary image function whose output is one if the visual stimulus is present at location (*x, y*) at the time of the *t*-th fMRI volume measurement, and zero otherwise.

The six parameters were fitted by maximum likelihood estimation, which was done by maximizing

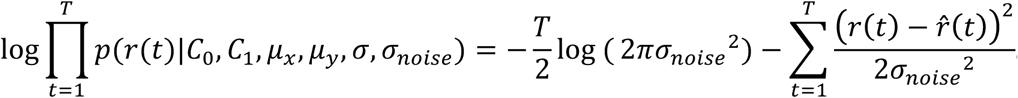

*T* is the number of the fMRI volumes used for model fitting, and we used 960 volumes from two experimental runs. *p*(*r*(*t*)*|C*_0_*, C*_1_*, μ_x_, μ_y_, σ, σ_noise_*) is the probability density function of *r*(*t*) given the six parameters. The maximization was conducted using a tool implemented in MATLAB (fminsearch.m from the optimization toolbox). To avoid local solutions, initial values in the optimization were searched on a regular grid. As per previous studies (Dumoulin and Wandell, 2008; Kay et al., 2008; Nishimoto et al., 2011), only well-fitted voxels were used in the analysis. From all available voxels (806±138, 922±85, 871±66, 664±164, 659±82, and 740±124 voxels for V1–V4, LOC, and FFA, respectively; mean±SD across subjects and sessions), we first eliminated the voxels whose estimated RF centers were outside the field the stimulus sphere could span (7.6° × 7.6°): 231±80, 255±55, 237±44, 183±65, 149±47, and 187±138 voxels for V1–V4, LOC, and FFA, respectively. Then we calculated the correlation coefficients between the real and fitted amplitudes to evaluate the fitness. The voxels with *r* > 0.2 were used (see Results for the numbers of selected voxels in individual areas).

The estimated models were also used in the decoding analysis. To separate data for decoding analysis and for RF model fitting, we performed a cross-validation procedure. In our experiments, each subject participated in the position tracking experiment that consisted of three experimental runs. Two runs were used for fitting receptive field models and the rest run was used as test data in the decoding analysis. The test run was shifted such that all runs were treated as test data once (leave-one-run-out cross-validation).

### Decoding analysis

We used the RF model or support vector regression (SVR)(Drucker et al., 1997; Chang and Lin, 2011) to predict the position of the stimulus from fMRI responses. In the prediction with the RF models, we calculated the stimulus position with the highest likelihood as the predicted position for each fMRI volume. Thus, the predicted position with the RF models was

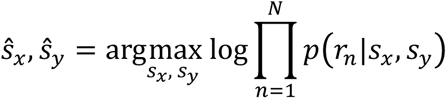

where

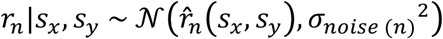

and

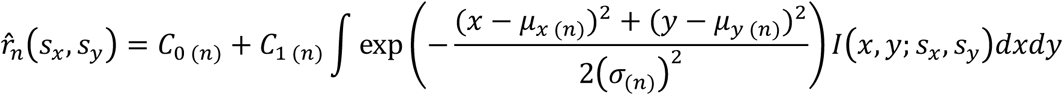

Here, *s*_*x*_ and *s*_*y*_ are the parameters that indicate the position of the stimulus center in the model, and *r*_*n*_ is the voxel amplitude of the n-th voxel in a given fMRI response. *p*(*r*_***n***_*|s*_***x***_, *s*_***y***_) is the probability density function of *r*_***n***_ given that the stimulus center is at (*s_x_*, *s*_***y***_). We assumed the Gaussian noise on different voxels to be independent, and the voxels in each visual area were combined by taking the product of their probability density functions. *C*_0_(*n*), *C*1(*n*)*, μ*_***x***_ (*n*), *μ_y_*(*n*), *σ*(*n*), and *σ_noise(*n*)_* are the RF model parameters for the *n*-th voxel. *I* (*x*, *y; s*_***x***_*, s_y_*) is the binary image function when the stimulus is centered on (*s_x_*, *s*_***y***_), thus the value of *I* (*x*, *y*; *s_x_*, *s_y_*) is one if the distance between (*x*,*y*) and (*s_x_, s_y_*) is less than the stimulus radius (0.8°), and zero otherwise.

For practical reasons, for each fMRI volume, we calculated the likelihood for each of 60 × 60 positions in the visual field and the position with the highest likelihood was treated as the predicted position.

In the prediction with SVR, the predicted position is given by

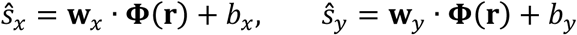

where

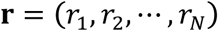

**w**_*x*_ and **w**_*y*_ are weight vectors, *b_x_* and *b_y_* are biases, and **Φ**(**r**) is a vector function that satisfies

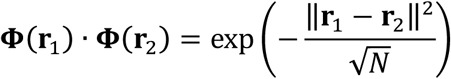

The models were trained by minimizing the cost function of the SVR algorithm with training data, and the model training and prediction were performed without explicitly calculating the weight vectors by using the kernel trick (Drucker et al., 1997; Chang and Lin, 2011) (RRID: SCR_010243).

We generated predicted positions for 1,440 fMRI volumes in three runs, and calculated the correlation coefficient between the true and predicted positions in the horizontal or vertical axes as the prediction accuracy.

## Results

First, we fitted an RF model to the response of each voxel (Dumoulin and Wandell, 2008). Our model consists of a two-dimensional Gaussian receptive field with the parameters of the mean (x, y positions) and the standard deviation (RF size). Gaussian noise is assumed in the response amplitude. To evaluate the fitness, we calculated the correlation coefficients between the real and fitted amplitudes (Figure 2A; *r* = 0.19±0.13, 0.20±0.13, 0.21±0.14, 0.18±0.12, 0.18±0.12, and 0.18±0.11 for V1–V4, LOC, and FFA, respectively; mean±SD across subjects and sessions). All visual areas analyzed show similar distributions of fitness. As per in previous studies, only well-fitted voxels with *r* > 0.2 were used in further analyses presented in the following (Dumoulin and Wandell, 2008; Kay et al., 2008; Nishimoto et al., 2011; 192±52, 237±47, 254±78, 135±84, 155±91, and 164±108 voxels for V1–V4, LOC, and FFA, respectively), while lower or higher thresholds on *r* yielded qualitatively similar results but with generally poorer decoding accuracies. Estimated RF sizes tended to be larger for voxels in the higher visual cortex (Figure 2B,C; ANOVA on mean RF sizes across visual areas, *F*(5,24) = 11.77, *p* = 7.968 × 10^−6^), consistent with previous studies (Dumoulin and Wandell, 2008; Amano et al., 2009).

**Figure 2.**
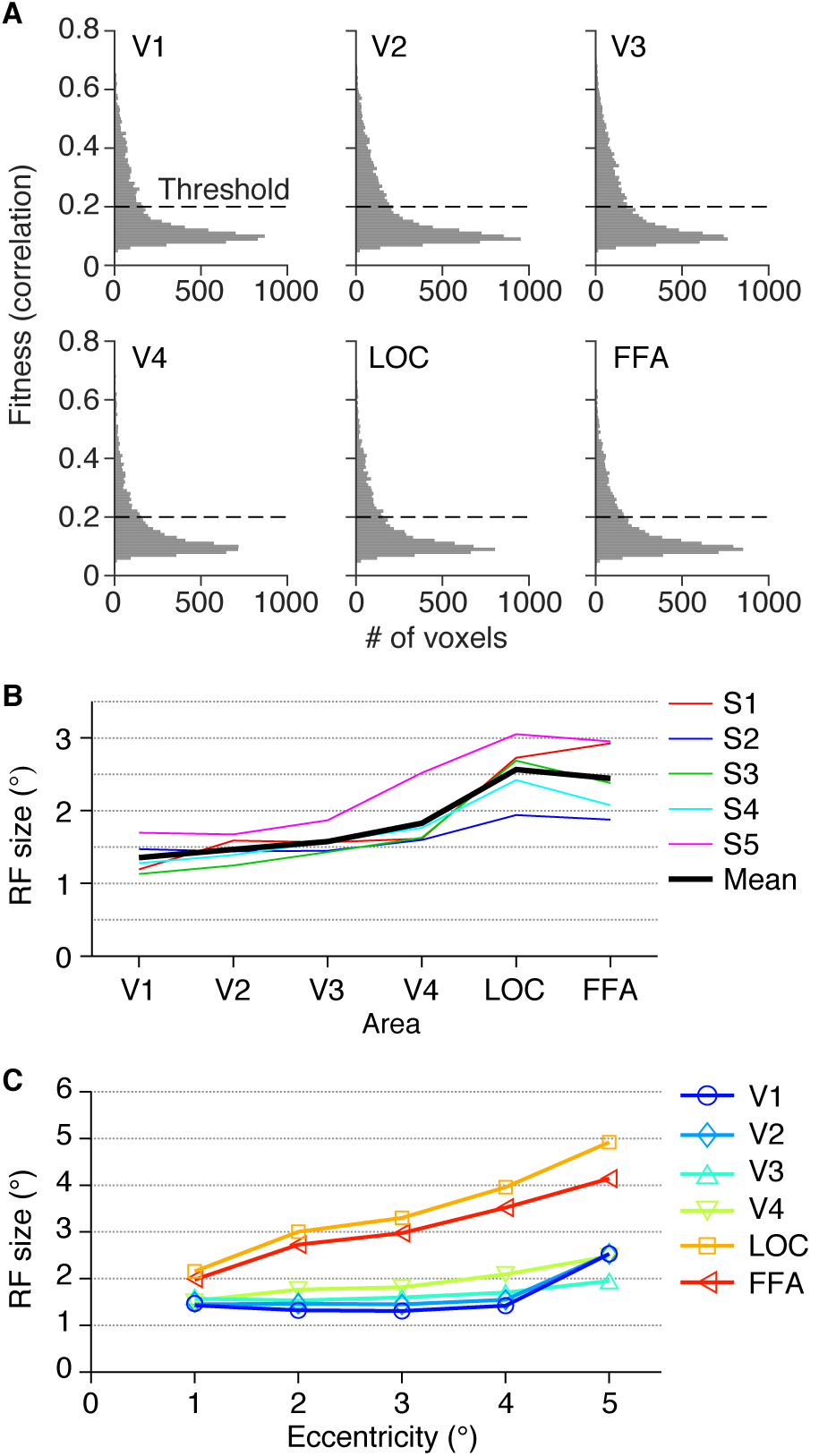
Properties of estimated receptive field models. **(A)** Histogram of the fitness of receptive field models. For each voxel, the correlation between the observed and fitted amplitudes was evaluated. Voxels were pooled across five subjects and three sessions. Voxels with estimated receptive field centers outside the field the stimulus could span were excluded. **(B)** Mean receptive field size for each visual area. We evaluated the receptive field size of each voxel using the parameter sigma of the fitted Gaussian receptive field. Colored lines show the mean across voxels for individual subjects. Black line shows the mean across subjects. **(C)** The relationship between eccentricity and receptive field size. The eccentricities of the estimated receptive field centers were binned into five levels with an interval of 1°. The mean receptive field size for each eccentricity level was calculated across voxels, and plotted as a function of the eccentricity.

Using the models described above, we conducted a decoding analysis to evaluate the amount of position information in each visual area. We estimated the 2D-coordinates of the ball position by taking the position with the highest likelihood for a given fMRI activity pattern. To quantify the prediction accuracy, we calculated the correlation coefficient between the true and predicted coordinates for each of the horizontal and vertical axes. Model fitting and position prediction were performed on fMRI data from separate runs via a cross-validation procedure (leave-one-run-out cross-validation).

The ball position was well predicted from the brain activity in all brain areas tested (Figure 3A,B upper; Movie 1): the correlation coefficients between the true and predicted positions (mean across subjects; horizontal/vertical coordinates) were 0.75/0.73 for V1, 0.74/0.74 for V2, 0.77/0.75 for V3, 0.63/0.62 for V4, 0.66/0.35 for LOC, and 0.66/0.40 for FFA (95% CIs: [0.54, 0.87]/[0.48, 0.87], [0.59, 0.84]/[0.53, 0.86], [0.50, 0.90]/[0.43, 0.90], [0.17, 0.86]/[0.21, 0.85], [0.47, 0.79]/[0.04, 0.60], and [0.50, 0.77]/[0.05, 0.66], respectively). Notably, the two higher visual areas with large RFs showed effective position decoding. All areas showed similar predictive performance for the horizontal position (Figure 3B upper, black line) with no significant difference (ANOVA on correlation coefficients [Fisher’s z-transformed] across visual areas, *F*(5,24) = 0.68, *p* = 0.6418). However, the decoding accuracy for the vertical position showed a decline in LOC and FFA (Figure 3B upper, gray line; *F*(5,24) = 3.27, *p* = 0.0216). In LOC and FFA, the decoding accuracy was significantly greater for the horizontal dimension than for the vertical dimension (*t*(4) = 11.98, *p* = 0.0003 for LOC; *t*(4) = 6.64, *p* = 0.0027 for FFA; *p* > 0.3 for all other areas). SVR yielded slightly greater decoding accuracies in general, with qualitatively similar dependencies on visual areas and the horizontal/vertical dimension (Figure 3B lower; ANOVA across visual areas, *F*(5,24) = 1.04, *p* = 0.4171 for the horizontal dimension, *F*(5,24) = 1.73, *p* = 0.1653 for the vertical dimension; *t*-test between the horizontal and vertical dimensions, *t*(4) = 4.09, *p* = 0.0150 for V4, *t*(4) = 8.06, *p* = 0.0013 for LOC, *t*(4) *=* 5.48*, p* = 0.0054 for FFA, *p* > 0.25 for all other areas), indicating that this tendency is independent of the decoding method. The difference found in LOC and FFA is consistent with the classification results in a previous fMRI study (Carlson et al., 2011), although the previous study did not test it for the lower visual cortex.

**Figure 3.**
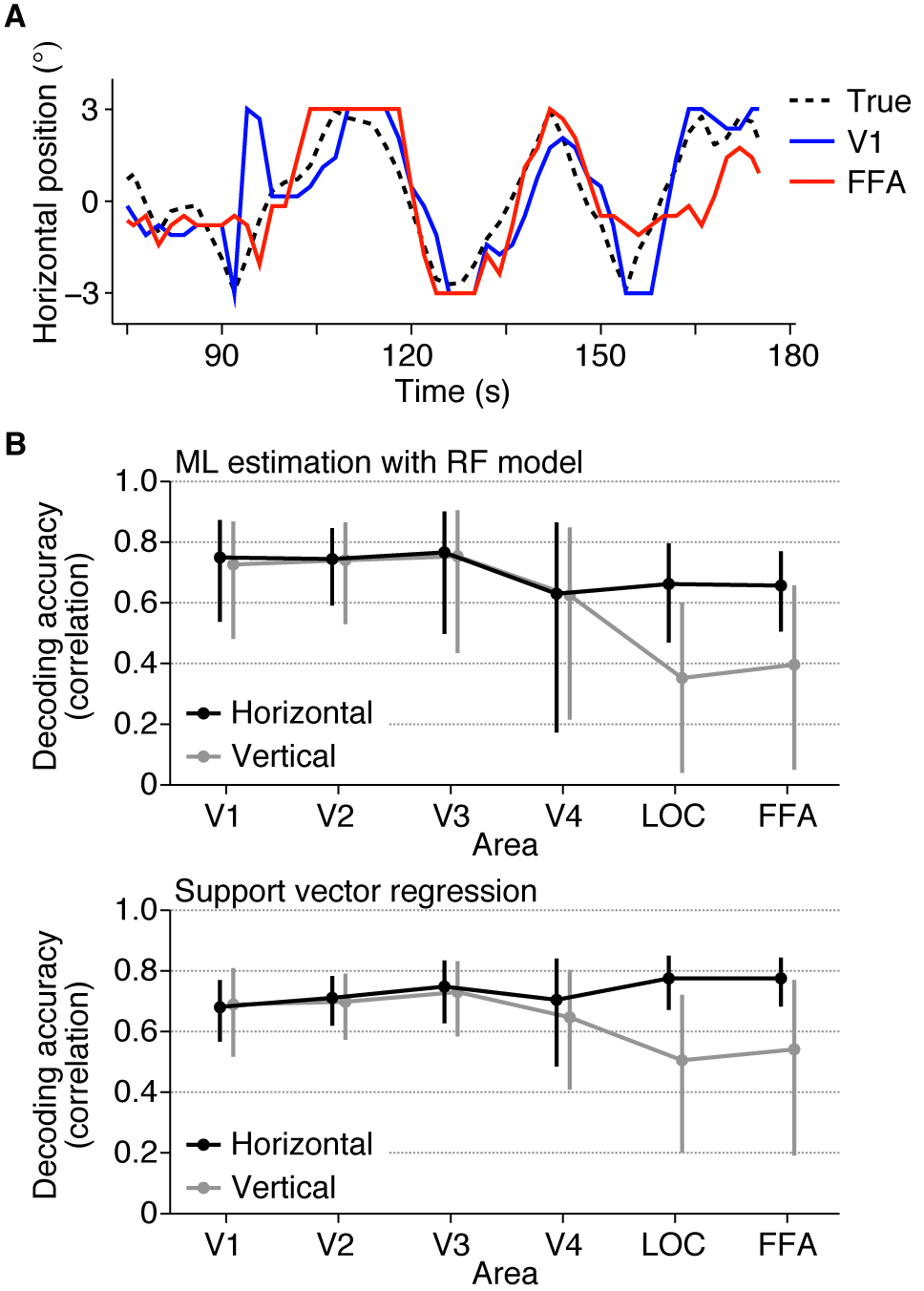
Position decoding accuracy from each visual area. **(A)** Examples of true and predicted trajectories of the ball position. The predicted trajectories were produced by maximum likelihood estimation using the receptive field models. **(B)** Decoding accuracy. The ball position was predicted from brain activity by maximum likelihood estimation with the RF models (upper) and SVR (lower). The accuracy was evaluated using the correlation coefficient between the true and predicted trajectories. The mean accuracies across subjects are shown. The calculations were performed separately for the horizontal (black line) and vertical (gray line) positions. Error bars show the 95% confidence intervals across subjects.

To find out factors that could affect the anisotropy in LOC and FFA, we examined the distribution of the RF centers of individual voxels in each area (Figure 4A,B). In LOC and FFA, the vertical positions of the RFs were narrowly distributed compared with V1–V4, while the horizontal positions of the RFs were distributed with similar degrees for all visual areas (ANOVA on the standard deviation of RF positions across visual areas, *F*(5,24) = 1.55, *p* = 0.2117 for the horizontal dimension, *F*(5,24) = 41.64, *p* = 4.635 × 10^−11^ for the vertical dimension; *t*-test between the horizontal and vertical dimensions, *t*(4) = 1.78, 7.82, 2.02, 3.27, 5.17, and 4.99, *p* = 0.1498, 0.0014, 0.1140, 0.0309, 0.0067, and 0.0076 for V1–V4, LOC, and FFA, respectively). This suggests that the lower decoding accuracies of LOC and FFA for the vertical dimension could be attributable to the narrow distribution of the RFs along this dimension, which is a factor not related to RF size.

**Figure 4.**
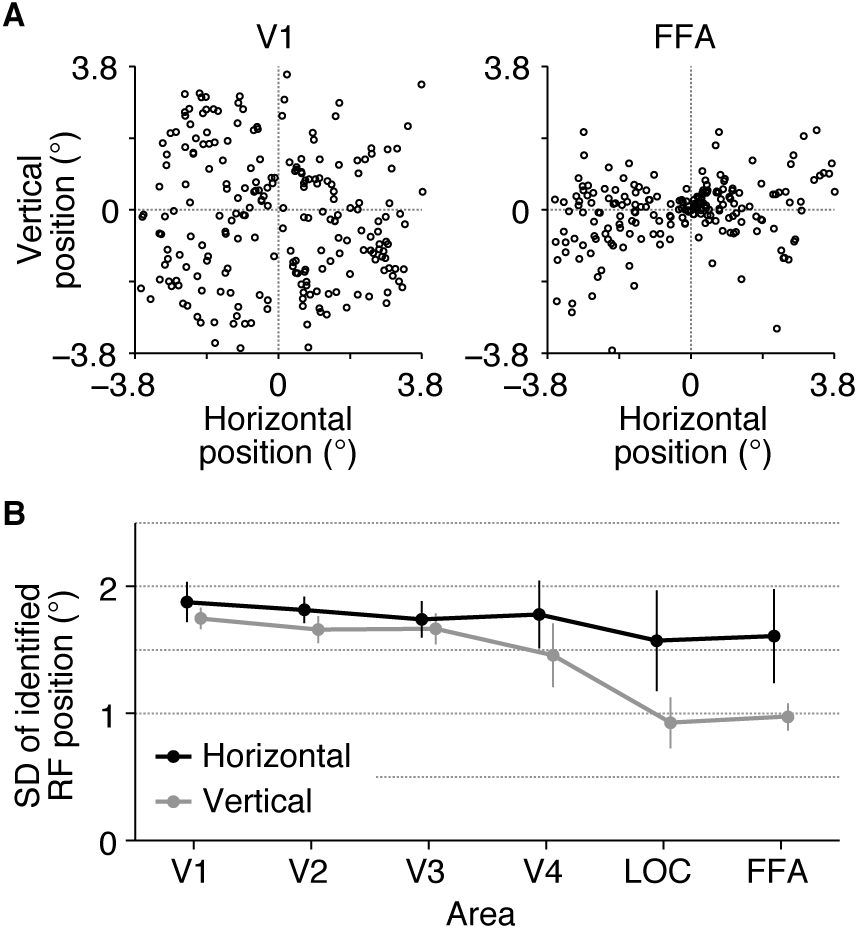
Spatial distribution of estimated receptive fields. **(A)** Examples of the distribution of estimated receptive field centers. Each circle shows the position of the receptive field center of a single voxel. We plotted the positions for the voxels in V1 and FFA from subject S3. **(B)** Standard deviation of the positions of receptive field centers. The mean values across subjects are shown. Error bars show the 95% confidence intervals across subjects.

The brain areas compared here contained different numbers of voxels. So, to confirm that the observed pattern of decoding performance across those visual areas was not due to the difference in the number of voxels used for prediction, we conducted the same decoding analysis with a fixed number of randomly selected voxels within each brain area. Similar comparison results were obtained independent of the number of voxels (Figure 5A,B).

**Figure 5.**
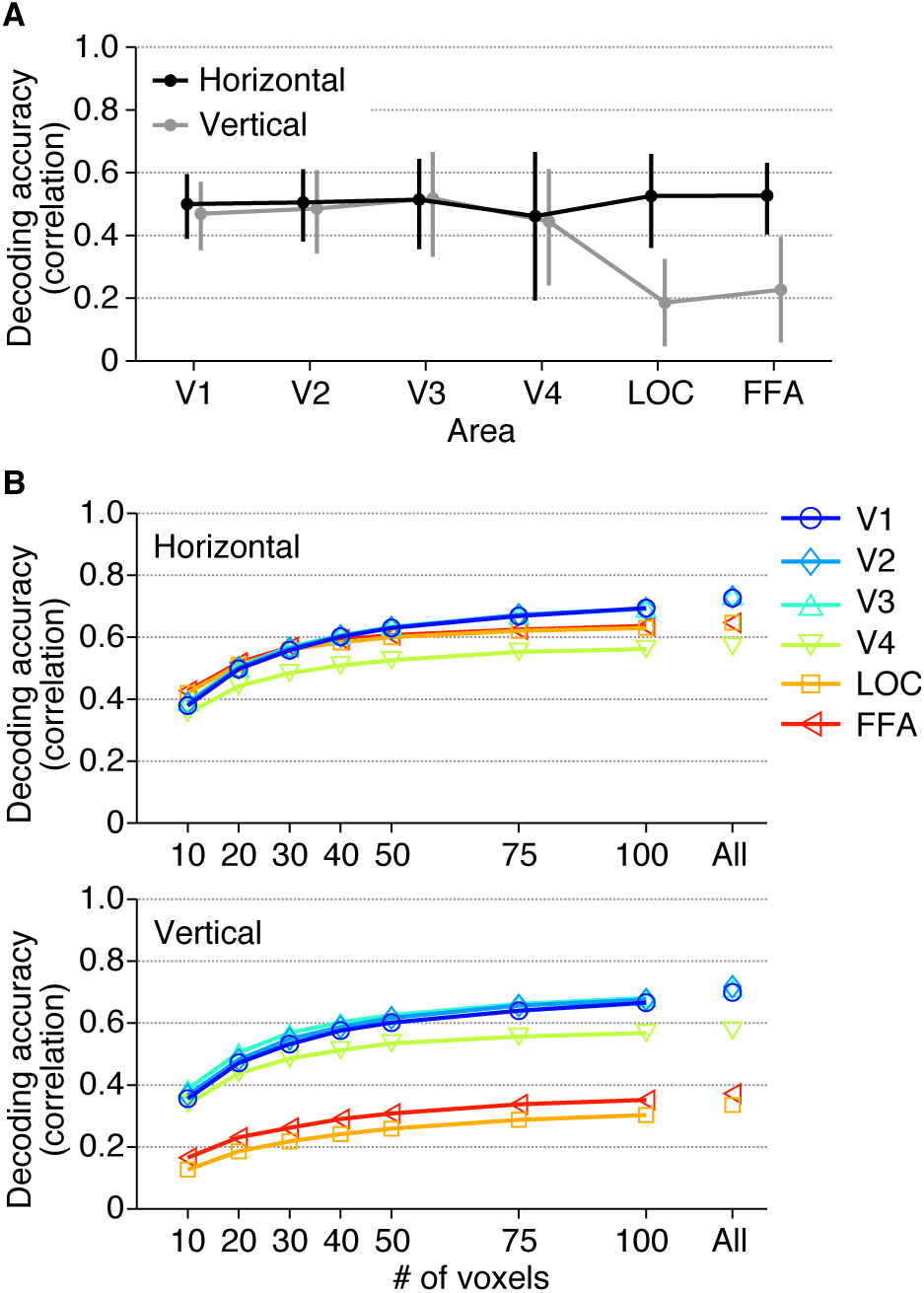
Decoding accuracy after matching the numbers of voxels. **(A)** Decoding accuracy with 20 voxels. The format is the same as in Figure 3B. We performed decoding analysis with RF models on brain activity from 20 randomly selected voxels in each visual area. Decoding accuracies were first averaged across 100 instances of random voxel selection in individual subjects, and then averaged across subjects. Error bars show the 95% confidence interval across subjects. After matching the numbers of voxels, we observed a similar tendency as in Figure 3B. This indicates that the tendency across visual areas was not caused by the difference in the number of voxels. **(B)** The relationship between decoding accuracy and the number of voxels. Decoding analysis was performed with a fixed number of randomly selected voxels with the same procedure. Mean decoding accuracies were plotted as functions of the number of used voxels.

Our results suggest that each visual area encodes position information similarly regardless of the difference in RF size if RF centers are equally distributed. However, it is possible that RF size can affect position coding when voxels are spatially restricted. We performed the position decoding analysis after excluding the voxels whose RF centers are near the stimulus position, while changing the threshold for the distance between a stimulus position and an RF center (Figure 6). To evaluate how steeply the decoding accuracy is degraded, an exponential decay function was fitted to the curve of the accuracy obtained from each visual area and subject. The resultant decay constant (*τ*) was used as a measure of the sensitivity to the distance threshold. We found that the decoding accuracies (tested on the horizontal dimension) for lower visual areas degraded more sharply than those for higher visual areas (ANOVA on decay constants across visual areas, *F*(5,24) = 5.47, *p* = 0.0017). Higher visual areas may be better at compensating for the loss of information with the large RFs of the remaining voxels. This observation suggests that RF size can be critical for encoding stimulus position with a limited neural population, although it is compatible with the fact that position encoding by the full population is equally accurate regardless of RF size.

**Figure 6.**
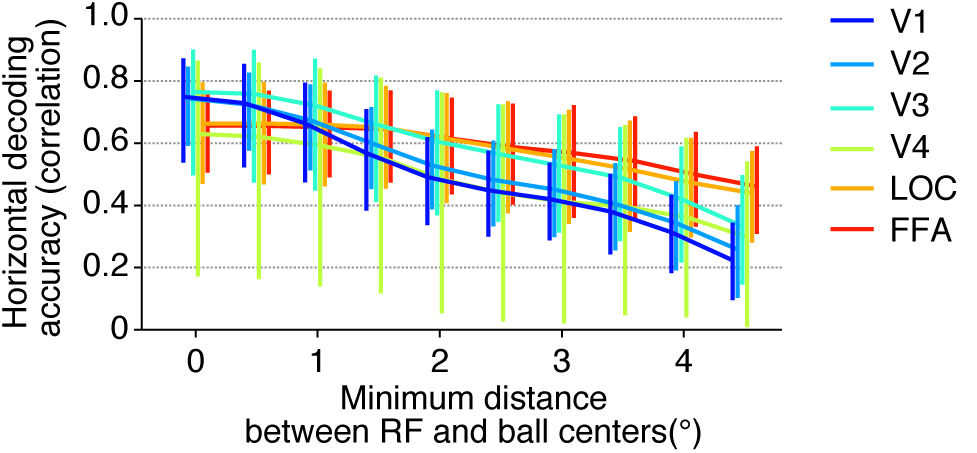
Decoding accuracy after excluding voxels whose receptive field centers are near the stimulus position. For each fMRI sample, we selected the receptive fields whose distances between the receptive field centers and the stimulus position are more than a threshold, and the stimulus position was predicted with those receptive fields by maximum likelihood estimation. The mean horizontal decoding accuracies across subjects for six visual areas are plotted as functions of the threshold. Error bars show the 95% confidence intervals across subjects.

## Discussion

In the present study, to investigate the relationship between the size of RFs and retrievable position information, we estimated RF sizes for fMRI voxels and evaluated how accurately the position of a seen object was predicted from activity patterns in each of six representative visual areas. We found that even with larger RF sizes, the position of the stimulus was predicted from activity patterns in high-level visual areas with similar accuracies to low-level visual areas especially for the horizontal position.

In the comparison of the decoding accuracy between the horizontal and vertical positions, the decoding accuracies for activity in LOC and FFA regarding the vertical position were lower than those for the horizontal position, and this anisotropy was not observed for the lower visual areas (Figure 3B). Although a previous fMRI study came to a similar conclusion on the anisotropy in LOC and FFA (Carlson et al., 2011), our study have compared lower to higher visual areas along the ventral cortical hierarchy using quantitative models. Furthermore, we demonstrated that these lower decoding accuracies are accompanied by a narrow spatial distribution of RFs for the corresponding direction (Figure 4), which may be a cause of the horizontal-vertical asymmetry in decoding accuracy. While similar decoding performance was obtained regardless of RF size in the condition where the centers of RFs used for prediction were distributed equally and widely, we also showed that when the voxel population was limited, RF size was critical for decoding accuracy (Figure 6). Thus, RF size may be important for spatial coding when a small neural population is used for inferring stimulus position. Further investigation of such collective properties of RFs would be useful for characterizing the mechanism and function of each brain region in representing position information.

Taken together, our findings provide experimental evidence that large RFs do not necessarily imply the loss of position information at the population revel. Regions in the higher visual cortex, such as LOC and FFA, appear to encode as much position information as the lower visual cortex, especially in the horizontal dimension, regardless of RF size. While our results demonstrate the availability of rich position information in higher visual cortex, it remains to be seen whether and how such information is used in later neural processing for recognition and behavior.

## Acknowledgements

The authors would like to thank Mitsuaki Tsukamoto, Makoto Takemiya, and Keiji Harada for helpful comments on the manuscript. This work was supported by Strategic International Cooperative Program (JST/AMED), ImPACT (Cabinet Office, Japan), and MEXT/JSPS KAKENHI grant no. 15H05920, 15H05710.

The authors declare no competing financial interests.

## Author contributions

YK and PS designed the study. PS and TH performed experiments. KM and PS performed analysis. KM, TH, and YK wrote the manuscript.

## Movie legend

**Movie 1.** Examples of true and predicted ball positions. The predicted positions were produced by maximum likelihood estimation using the RF models.

